# Comparative microbiome analyses reveal differences between wild populations and captive groups of the Montseny Brook Newt (*Calotriton arnoldi*)

**DOI:** 10.1101/2025.06.18.660306

**Authors:** Sergi A. Tulloch Jiménez, Maria Estarellas, Dean C. Adams, Anthony Bonacolta, Viviana Pagone, Daniel Fernández-Guiberteau, Fèlix Amat, Albert Montori, Francesc Carbonell, Elena Obon, Mónica Alonso, Marta Santmartín, Josep Xarles, Rosa Marsol, Daniel Guinart, Sònia Solórzano, Adrián Talavera, Bernat Burriel-Carranza, Elena Bosch, Javier del Campo, Salvador Carranza

**Affiliations:** Institut de Biologia Evolutiva (CSIC-Universitat Pompeu Fabra), Pg. Marítim de la Barceloneta, 37 49, 08003 Barcelona, Catalonia, Spain; Department of Ecology, Evolution and Organismal Biology, Iowa State University, USA; Department of Marine Biology and Ecology, Rosenstiel School of Marine, Atmospheric and Earth Science, University of Miami, Miami, Florida, 33149 USA; CREAC - Centre de Recerca i Educació Ambiental de Calafell (GRENP-Ajuntament de Calafell), Spain; Àrea d’Herpetologia, BiBIO, Museu de Granollers – Ciències Naturals. Palaudàries 102, Granollers 08402, Spain; Àrea de Gestió Ambiental Servei de Fauna i Flora (Centre de Fauna de Torreferrussa), Spain; Zoo de Barcelona, Spain; Àrea de Gestió Ambiental Servei de Fauna i Flora (Centre de Fauna del Pont de Suert), Spain; Servei de Gestió de Parcs Naturals, Diputació de Barcelona, Spain; Museu de Ciències Naturals de Barcelona, P° Picasso s/n, Parc Ciutadella, 08003, Barcelona, Spain; Institute of Evolutionary Biology (UPF-CSIC), Department of Medicine and Life Sciences, Universitat Pompeu Fabra, Barcelona 08003, Spain

**Keywords:** microbiome, captivity, amphibians, conservation, reintroduction, newts

## Abstract

The Montseny brook newt, *Calotriton arnoldi*, is a Critically Endangered amphibian species endemic to the Montseny Massif in Catalonia, Northeastern Spain. Due to population declines and threats to its natural habitat, an *ex-situ* breeding program was initiated in 2007. A key goal of the program is to ensure the survival of captive-bred individuals after reintroduction, which in amphibians heavily relies on the specimens’ microbiome being capable of protecting them from environmental microorganisms, especially considering the global Chytridiomycosis pandemic caused by the fungi *Batrachochytrium dendrobatidis* (*Bd*) and *Batrachochytrium salamandrivorans* (*Bsal*). This study aims to characterize the microbiome of wild and captive specimens of *Calotriton arnoldi*, to identify differences in microbiome composition, and to determine their potential impact on captive-bred individuals upon reintroduction. Up to 7,438 ASVs (Amplicon Sequence Variants) were identified from 138 samples from 21 and 61 wild and captive-bred individuals, respectively. Results indicate that wild populations from different subspecies have significantly different microbiome composition, as do wild and captive-bred groups from the same subspecies.

Additionally, dissimilarities in microbiome variability were only found within each subspecies, between wild and captive-bred groups. In terms of composition, certain bacteria were identified as potential markers for both wild and captive environments. Enhancing microbiome variability might improve the survival prospects of reintroduced specimens. Thus, exposing captive specimens to a more natural environment while in captivity or a soft-release procedure could potentially mitigate the absence of exposure to other bacteria and potential pathogens from their native environment.

## INTRODUCTION

Recent studies based on the analysis of 32% of terrestrial vertebrate species indicate that beyond the ongoing global extinctions, our planet is undergoing a rapid decline and disappearance of natural populations, referred to as “biological annihilation” [1]. Over the past decades, factors such as overexploitation, habitat loss, the introduction of invasive species, pollution, climate change, and emerging diseases have led to a catastrophic decline in the number and size of vertebrate species populations [2, 3]. As a result, in the last 100 years, hundreds of species and vertebrate populations have become extinct at a rate 100 times higher than the natural extinction rate over the past two million years, suggesting we are already in the sixth major episode of mass extinction on our planet [1, 3].

Among all groups of terrestrial vertebrates, amphibians have received significant attention in the last four decades. Despite having survived several mass extinctions, evidence indicates that more amphibian species have already gone extinct or are endangered compared to other vertebrate groups [3]. A reason for such increased susceptibility is their permeable skin, allowing the absorption of water and gases for respiration and hydration but, at the same time, making them especially susceptible to pollution, climate change-related events (such as changes in temperature or precipitation patterns), and emerging diseases.

One of the most significant challenges amphibians face is the chytridiomycosis pandemic, caused by the chytrid fungi *Batrachochytrium dendrobatidis* (*Bd*) and *Batrachochytrium salamandrivorans* (*Bsal*), and viral infections (e.g. *Ranavirus*), already leading to the disappearance of over 200 amphibian species worldwide with many more predicted to become extinct in the near future [4–5]. Chytrids are fungi that usually live in soil or water but occasionally parasitize other fungi, plants or insects. Importantly, *Bd* and *Bsal* are the only known chytrids that infect vertebrates. These species remain as spores in the water until they encounter a host. At this point, they excyst, fructify, and proliferate throughout the host’s keratinized body parts (i.e. mouthparts in larval stages and skin in adults) [6], disrupting the normal regulatory functioning of the amphibians’ skin [7].

Because of its permeability, the amphibian skin has a very important microbial component based on bacteria, fungi, and protists; yet how they obtain their microbiota remains unclear. Evidence suggests that individuals may acquire their microbiome from the environment, both through horizontal transmission (e.g., during mating or communal gatherings) [8] and vertical transmission (particularly in species that exhibit parental care) [9]. The microbiome has a major influence on many processes, including the host’s digestion, behaviour, development, and reproduction [10–12], but what is of most interest for this study is the pivotal role it has as an integral part of the immune system [13]. Harris et al. [14] demonstrated that some skin bacteria inhibit the growth of chytrid fungi, emphasizing the importance of the individual’s microbial community and its impact on disease survival. Moreover, it has been shown that some amphibians can enhance their chances of survival against chytrid fungi if they have been previously exposed to a milder strain of the fungus [15].

The Montseny brook newt, *Calotriton arnoldi*, is a species endemic to the Montseny Massif in eastern Catalonia, formally described by Carranza and Amat in 2005 [16]. Its area occupancy is limited to eight streams in less than 10 km^2^, making it the most threatened amphibian species in Europe and being considered Critically Endangered by the IUCN [17]. Bearing in mind that *Calotriton arnoldi* is a completely aquatic urodele at both larval and adult stages, it faces constant threats, including water overexploitation, deforestation, stream continuity disruption, warming temperatures, natural disasters, and emerging diseases like chytridiomycosis. This is particularly concerning given that a recent *Bd* and *Bsal* outbreak was detected in the Montnegre i el Corredor Natural Park, located just 15 km south of the natural range of *C. arnoldi* [18].

Recently, two subspecies of *Calotriton arnoldi* have been recognized: *C. a. laietanus*, comprising five populations located to the west of the Tordera River (Western populations: B1–B5), and *C. a. arnoldi*, consisting of three populations to the east (Eastern populations: A1–A3) [16, 19]. Its census is also worryingly low, indicating that only 1,000-1,500 individuals remain in their natural habitats [16]. In response to the critical conservation status of the species, an *ex-situ* breeding program was initiated in 2007, followed by the launch of a LIFE project in 2016 (LIFE15 NAT/ES/000757) aimed at improving the species’ chances of survival. The initial breeding center, the Torreferrussa Wildlife Recovery Centre, housed founding individuals from both subspecies: individuals from the Western populations B1 and B2 (*C. a. laietanus*) and from the Eastern A1 population (*C. a. arnoldi*), which were kept in separate enclosures, as the subspecies of *C. arnoldi* in the wild are separated by a natural barrier. Six years later, the Barcelona Zoo joined the program. Unlike the original center, however, the founding individuals at the Barcelona Zoo consisted of first-generation newts bred at Torreferrussa for both subspecies. In 2013, the breeding program was further expanded with the inclusion of a third facility, the Pont de Suert Wildlife Recovery Centre, which also began with first-generation *C. a. laietanus* individuals from Torreferrussa. Although *ex-situ* breeding programs are are valuable tools for species recovery, they also present certain drawbacks. These include potential genetic risks, such as adaptation to captivity– where traits favored in captive conditions may reduce fitness in the wild–as well as disparities in microbiomes between wild and captive populations, or loss of genetic diversity [20–22]. These concerns are particularly relevant in this case, as some genetic clusters are not currently represented in the breeding program [23].

This study aims to characterize the microbiome of *Calotriton arnoldi* across both recognized subspecies, examining individuals from both wild and captive populations. We also seek to identify and compare differences in microbiome composition between these environments. Additionally, we discuss the potential role of the microbiome in influencing the species’ survival, both under captive conditions and following reintroduction into the wild.

## MATERIALS AND METHODS

### Sampling and sampling sites

A total of 138 microbiome skin samples of *Calotriton arnoldi* were collected during May 2022 and between March and April 2023. 29 samples were obtained from 21 individuals of the *C. a. laietanus* population B1, and 10 samples from 5 individuals of the *C. a. arnoldi* population A1. *Ex-situ* samples were collected from all three breeding centers, with attention to subspecies and generation. From the Torreferrussa Wildlife Recovery Centre a total of 27 specimens were sampled, including both subspecies and all three available generations (F0, F1, and F2), to obtain 20 samples per subspecies (Fig. 1). From the Barcelona Zoo, 28 samples were collected from 23 individuals belonging to both subspecies and two available generations (F1 and F2) (Fig. 1). Finally, at the Pont de Suert Wildlife Centre, 19 samples were collected from 11 individuals of *C. a. laietanus*, representing both available generations (F1 and F2) (Fig. 1).

**Fig. 1:**
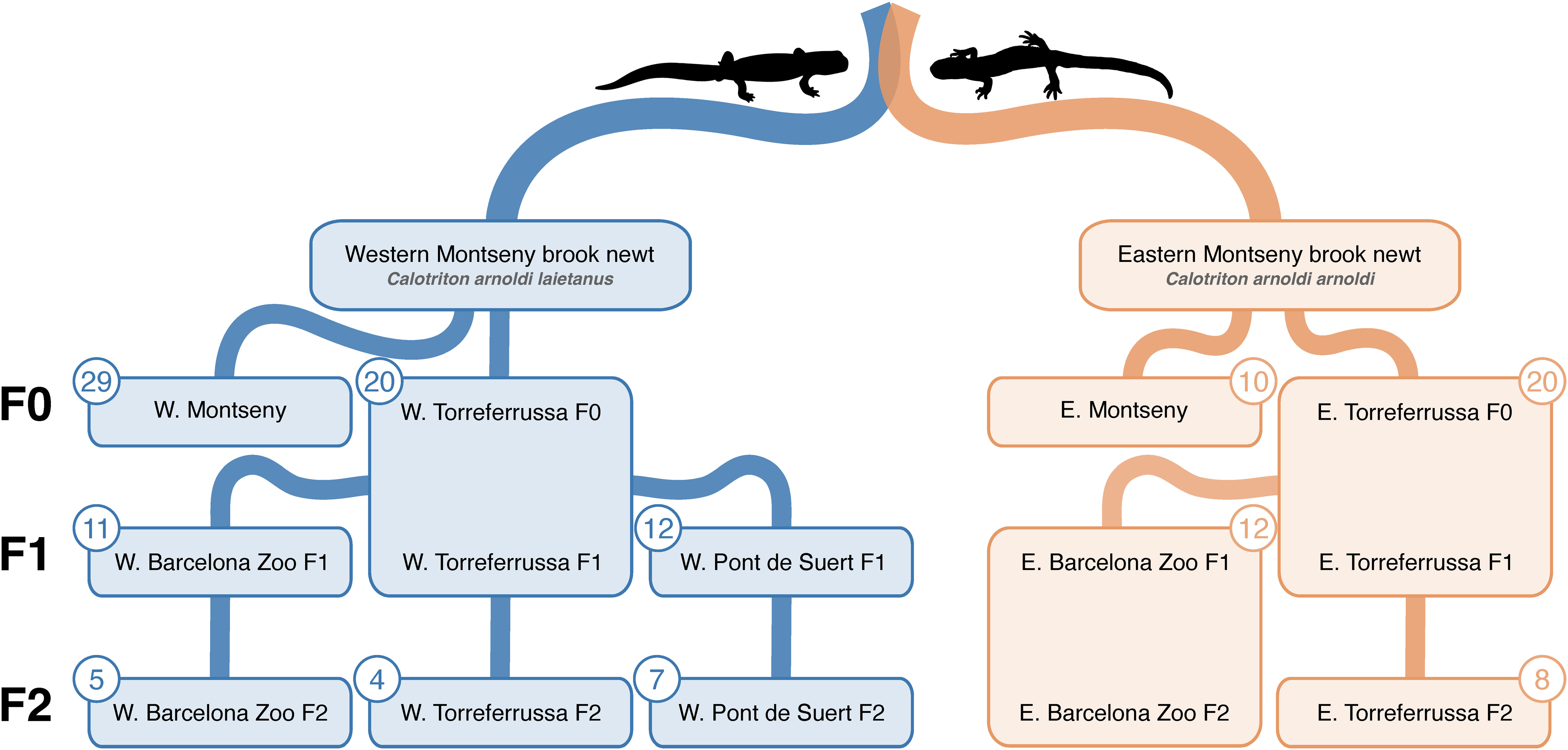
Summary of sample sorting, including subspecies, location, generation, assigned group and amount of skin microbiome samples per group. All group names were simplified according to their subspecies (W: Western Montseny brook newt (*C. a. laietanus*); E: Eastern Montseny brook newt (*C. a. arnoldi*)), location, and the generations they contain (F0, F1 and/or F2). Silhouettes were extracted from images by A. Talavera.

Water samples were also collected using swabs from each group of *C. arnoldi.* Groups of individuals were considered isolated when housed in water systems that did not mix at any point. Based on this criterion, all individuals were classified into eleven distinct groups (Fig. 1), which serve as the basis for most of the statistical analyses conducted in this study.

To minimize sex-based sampling bias, individuals were sexed during swab collection whenever possible. Two skin swabs were taken from each sampled individual following Bletz et al. [24]. Sampling was conducted according to the sanitary protocol of the Generalitat de Catalunya [25] to avoid contamination and the spread of emerging diseases. Permits to carry out this work were granted by the Wildlife Service of the Generalitat de Catalunya and the Area of Natural Parks of the Diputació de Barcelona.

### DNA extraction from swabs and sequencing

DNA was extracted from both swabs using the DNeasy Blood and Tissue Extraction kit (QIAGEN) following a slightly modified protocol for skin bacterial DNA [24]. The V4 region of the bacterial 16S ribosomal RNA gene was amplified with a PCR following an adapted protocol developed by the Earth Microbiome Project [26]. The PCR reaction per sample included 0.5 μL of both forward (515 F: 5’-TCG TCG GCA GCG TCA GAT GTG TAT AAG AGA CAG GTG YCA GCM GCC GCG GTA A-3’) and reverse (806 R: 5’-GTC TCG TGG GCT CGG AGA TGT GTA TAA GAG ACA GGG ACTACN VGG GTW TCT AAT-3’) primer and adapter (10 μM;), 22.5 μL of Taq SuperMix (Invitrogen Platinum PCR SuperMix, High Fidelity) and 3 μL of the extracted DNA. PCR conditions were as follows: denaturalization at 94°C for 3 minutes, followed by 30 cycles of 30 seconds at 94°C, 30 seconds at 52°C and 30 seconds at 68°C, with a final extension step at 68°C for 4 minutes. Subsequently, PCR products were visualized in a 1% agarose gel. Finally, PCR products were sequenced for an average of 50 000 reads per sample using paired-end 2×250 v2 chemistry on Illumina MiSeq at the Genomics Core Facility of the Pompeu Fabra University (Barcelona Biomedical Research Park, Barcelona, Spain). The DNA gene amplicon reads will be deposited in the NCBI Sequence Read Archive (PRJNA1265703).

### Microbiome Analyses

All statistical analyses were performed in R v.4.3.1 [27] and are available upon request. Each read was trimmed of its primers and sequencing adapters using Cutadapt [28]. Then, DADA2 v.1.28.0 [29] was used to assess read errors, truncate reads, and merge paired-end reads, followed by chimera removal. Afterwards, Amplicon Sequence Variants (ASVs) were inferred using a Bayesian classifier with the Silva database [30]. The Phyloseq v.1.44.0 package [31] was used to manipulate the amplicon sequence data within R, as well as to assess alpha diversity and generate relative abundance plots. Lab/kit contaminants were removed from the ASV table using the R package DECONTAM v.1.20.0 [32]. Chloroplast, mitochondrial, eukaryotic, and embryophyte ASVs were also removed. Following this quality control step, the final dataset comprised 7 742 406 processed reads assigned to 7,438 unique ASVs (<u>Table S1</u>). The dataset included 138 samples from 82 *C. arnoldi* individuals (Fig. 1) and 11 water samples.

For the statistical analyses, a centre-log-ratio transformation of the dataset was first performed using CoDaSeq v.0.99.6 [33] to account for the compositional nature of the data. Compositional and variance differences were tested for statistical significance using the RRPP v.1.3.1 package [34] linear models (with 10 000 permutations), in which permutational ANOVA was performed that accounted for both *location* (each breeding center or the wild) and *group*. When positive correlations were found, pairwise comparisons were conducted to explore the data further. Multiple comparisons were evaluated adjusted using Bonferroni correction to control for experimentwise type I error. Next, alpha diversity was assessed through the Shannon Diversity Index [35] and the Chao1 Index [36]. Ampvis2 v.2.8 [37] was used to generate heatmaps and identify core ASVs between sample metadata. Core ASVs were defined as those shared by at least 75% of the samples with a relative abundance equal or greater than 0.1%. ANCOMBC v.2.2.0 [38] was used to determine significantly different relatively abundant ASVs between groups, considered significant when (|Log_2_ (FC)|) > 2 and p-value < 0.05. BLAST [39] was used to identify relevant ASVs when the Silva database taxonomy was not sufficiently specific.

## RESULTS

### Diversity analysis

To compare the microbiomes of the two *Calotriton arnoldi* subspecies, only samples from wild individuals were included in the analysis. This approach allowed us to minimize the potential confounding effects of captivity and focus specifically on differences between the Western and Eastern Montseny brook newts. No significant differences were found when comparing their alpha diversity through the Shannon Diversity Index and the Chao1 Index (Fig. 2A & 2B). Still, the subsequent permutational ANOVA analysis revealed significant differences in microbiome composition (Z = 2.5735, p = 0.0002) but not in the variance of the microbiome (*C. a. laietanus* = 16 461.39, *C. a. arnoldi* = 13 555.42, *p* = 0.2757) (Fig. 2C).

**Fig. 2:**
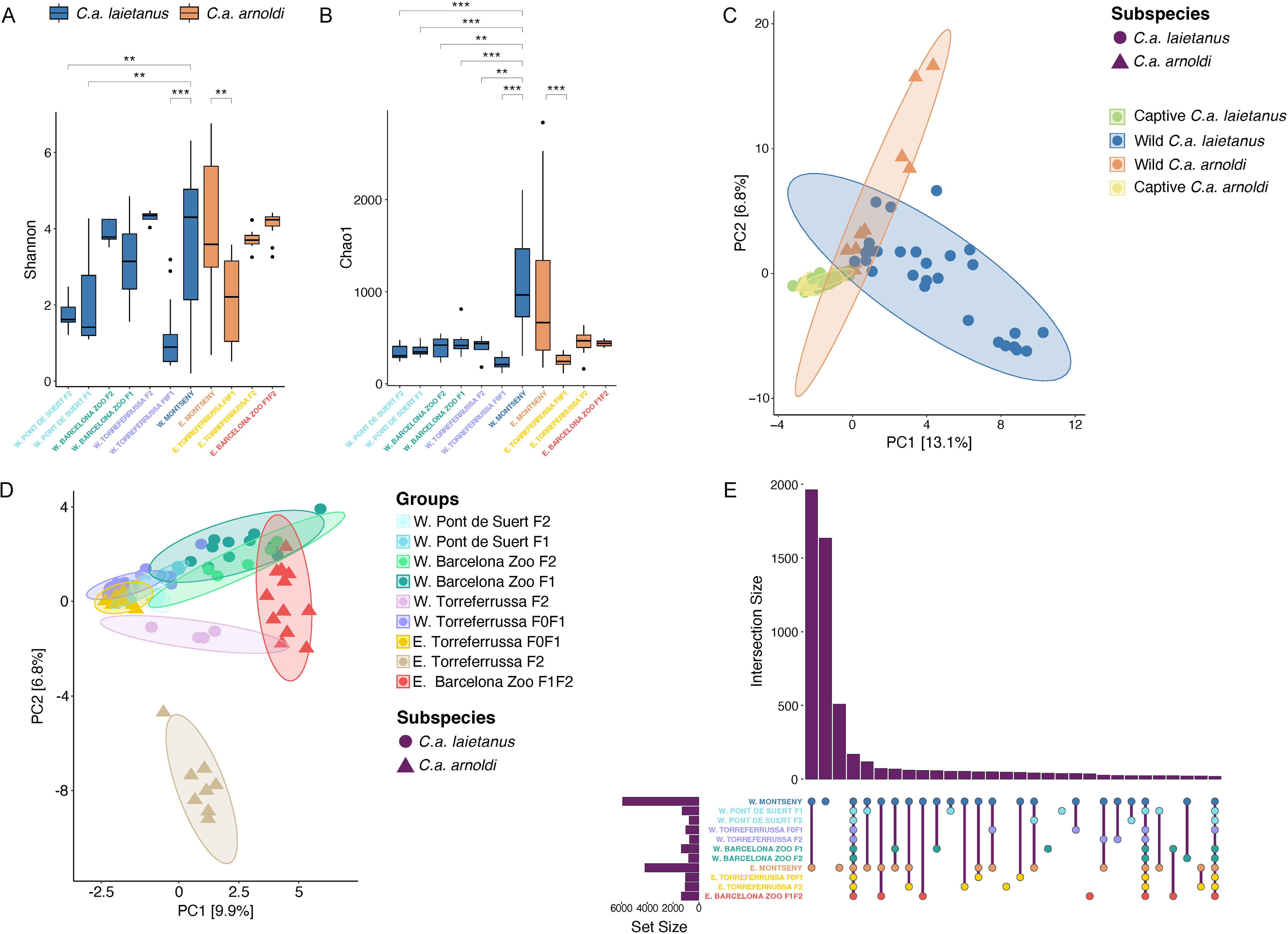
Alpha diversity, composition and variance differences and shared ASVs across wild populations and captive groups of *C. arnoldi*. Wild and captive-bred comparison of **A** Shannon’s Diversity Index and **B** Chao1 Index, with P-value for pairwise comparison indicated above (0.05 ‘*’ 0.01 ‘**’ 0.001 ‘***’ 0). Aitchison Distance PCA with 95% confidence interval ellipse for **C** Wild and captive-bred samples separated by subspecies and **D** Captive-bred groups. **E** UpSet diagram for shared ASVs across all groups of *C. arnoldi.* W: Western Montseny brook newt (*C. a. laietanus*); E: Eastern Montseny brook newt (*C. a. arnoldi*).

Similarly, all captive-bred groups were compared to the wild groups from their same subspecies (Fig. 2A & 2B). The Shannon Diversity Index displayed significant differences between the wild *C. a. laietanus* and three captivity groups: W. Torreferrussa F0F1 and both groups from Pont de Suert (F1 and F2). These wild-captivity differences were much more accentuated in the Chao1 Index comparison, where all captive *C. a. laietanus* groups significantly differed from the wild one. As for the Eastern Montseny brook newt, only E. Torreferrussa F0F1 significantly differed from the wild group in both alpha diversity indexes (Fig. 2A & 2B).

The permutational ANOVA performed for each subspecies revealed that both *location* and *group* variables were significant in the model (for *location*, Z = 5.8451 in *C. a. laietanus* and Z = 4.0966 *C. a. arnoldi*, with *p* < 0.0001 in both subspecies. As for *groups,* Z = 1.9052 with *p* = 0.0264 in the Western Montseny brook newt and Z = 6.4011 with *p* < 0.0001 in the Eastern Montseny brook newt). Subsequently, pairwise comparisons (<u>Table S2</u>) found that microbiome composition did not differ between groups from the same location (except for the E. Torreferrussa F0F1 and F2 groups) but did differ when compared to their wild counterpart. The variance displayed the same behavior, although in this case without exceptions (Fig. 2C & 2D). Reinforcing this idea, the wild populations were found to share more ASVs between their microbiomes than captive*-*bred groups presented in their whole microbiome (Fig. 2E).

### Compositional analysis

*C. arnoldi*’s microbiome mainly comprises two phyla: Proteobacteria and Bacteroidota. Most of the Proteobacteria belong to Gammaproteobacteria, including the most abundant genera, which can dominate an entire group, like *Acinetobacter* in W. Torreferrussa F0F1 or *Nevskia* sp. in both Pont de Suert F1 and F2 groups (Fig. 3A & 3B). Interestingly, all populations/groups share *Streptococcus* ASVs and an *Oceanotoga* sp.; most populations/groups also have *Pseudomonas* spp. Some genera are only significantly present in the wild populations, such as an unclassified species from the genera *Cytophagales* or *Verrucomicrobiales.* On the other hand, captive-bred groups seem to share *Flavobacterium* ASVs, specifically ASV 7, 12, and 16 (Fig. 3B). Moreover, the relative abundances of ASVs in *C. arnoldi*’s skin seemed to be unrelated to the relative abundances of ASVs found in the water (<u>Fig. S1</u> and <u>Table S3</u>).

**Fig. 3:**
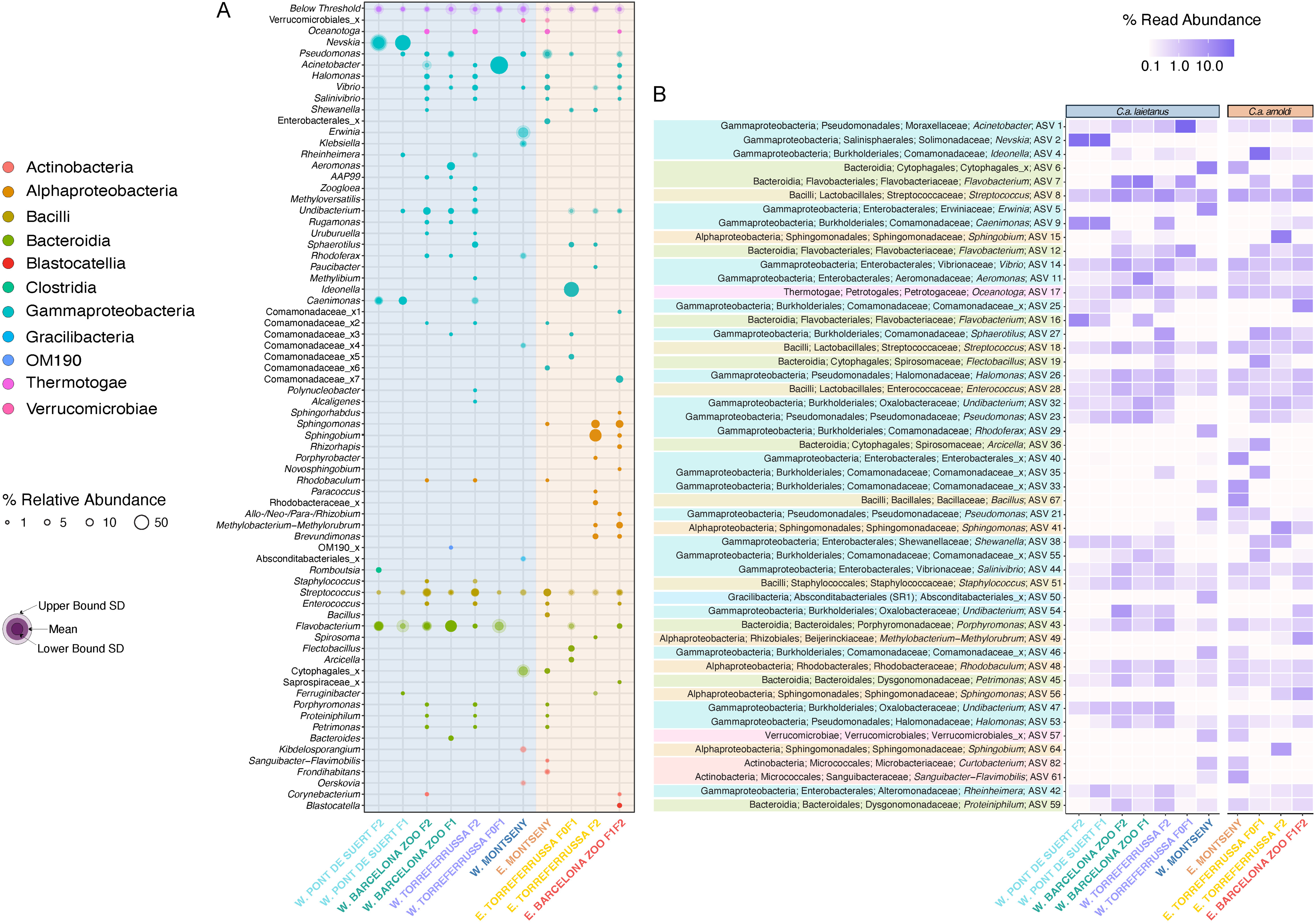
Microbiome composition of wild populations and captive-bred groups of *C. arnoldi*. **A** Bubble plot representing Genera with >1% relative abundance in at least one of the wild populations and captive-bred groups and **B** Heatmap representing the most abundant ASVs across all wild populations and captive-bred groups; both color-coded according to taxonomic Class. W: Western Montseny brook newt (*C. a. laietanus*); E: Eastern Montseny brook newt (*C. a. arnoldi*).

### Core microbiome

The core microbiome of *C. arnoldi*, defined as the set of ASVs shared across all samples, consisted of 20 ASVs, with ASV 8 standing out as the most prevalent in larger abundances (Fig. 4A). However, when analyzing the core microbiomes of each wild population and captive-bred group, no ASVs were consistently shared across all populations and groups (Fig. 4B). When excluding the *C. a. laietanus* group from Torreferrussa (W. Torreferrussa F0F1), which was dominated by ASV 1 with a relative abundance of 78.94% (Table S3), two previously mentioned genera emerged as core bacteria: *Streptococcus varani* (ASV 8) and *Oceanotoga* sp. (ASV 17). Additionally, eight core ASVs were identified across the wild populations (Table S4), including the two above, as well as *Vibrio* sp. (ASV 14) and another *Streptococcus* sp. (ASV 18). No core ASVs were shared across all captive-bred groups. However, *Undibacterium* sp. (ASV 47) appeared as a distinguishing feature in *C. a. laietanus* captive-bred groups (excluding W. Torreferrussa F0F1), while a different *Undibacterium* sp. (ASV 32) and *Sphaerotilus* sp. (ASV 27) were found to characterize the *C. a. arnoldi* captive-bred groups (Table S4).

**Fig. 4:**
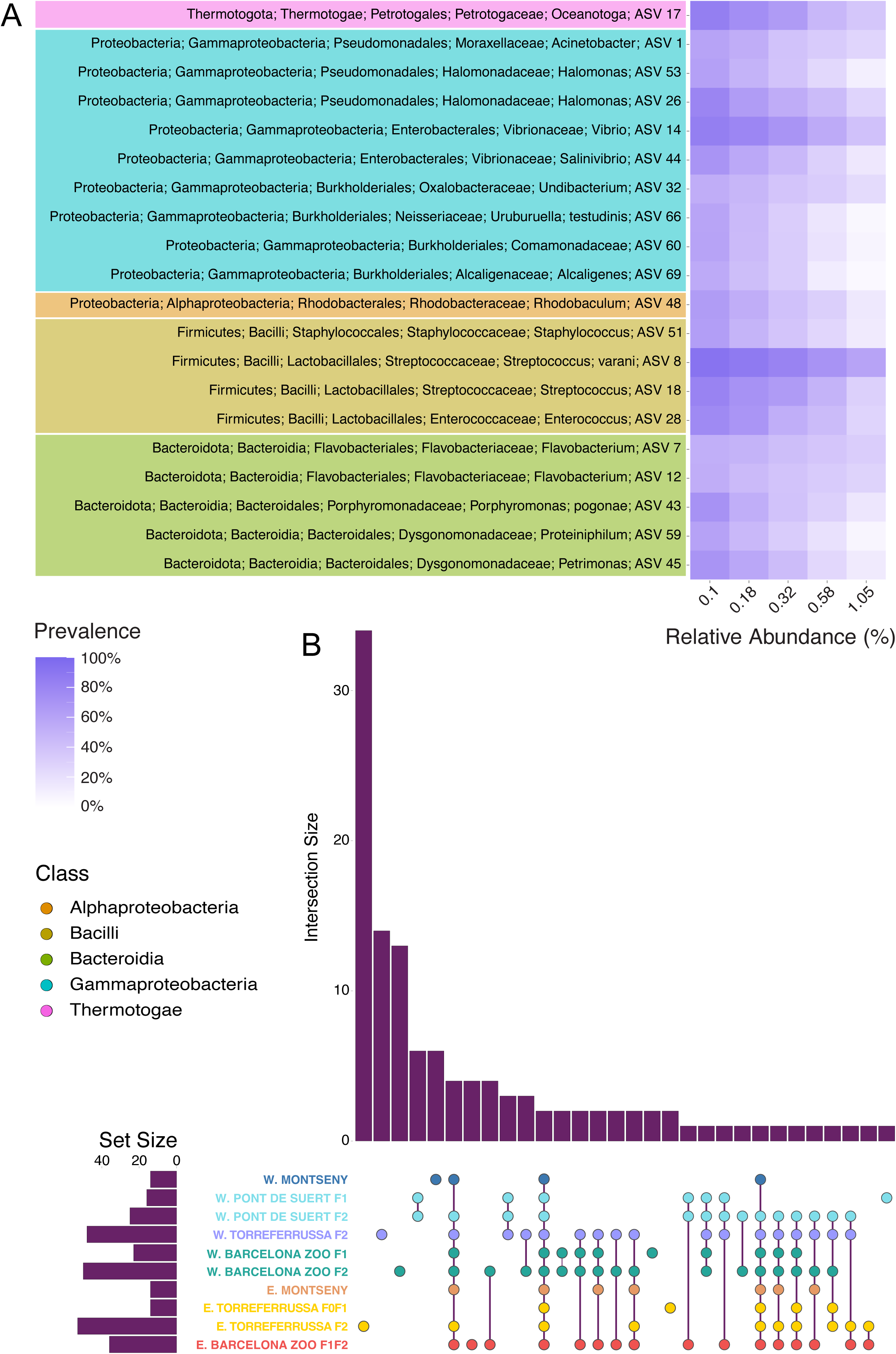
Core bacteria of *C. arnoldi’*s microbiome. **A** Heatmap representing core ASVs for all samples pooled together, coloured according to taxonomic Class. **B** UpSet diagram for shared core ASVs across wild populations and captive-bred groups. W: Western Montseny brook newt (*C. a. laietanus*); E: Eastern Montseny brook newt (*C. a. arnoldi*).

### Differential abundance

To further explore microbiome differences, ANCOM-BC analyses were conducted on selected pairwise comparisons involving both wild populations and captive-bred groups (<u>Table S5</u>). None of the core ASVs identified in the wild populations showed significant differential relative abundance when comparing the two subspecies. However, 101 ASVs were significantly more abundant in the wild Western Montseny brook newt (*C. a. laietanus*), including ASV 6, ASV 29, and ASV 50 -all of which are core ASVs for this population. In contrast, 47 ASVs were significantly more abundant in the wild Eastern Montseny brook newt (*C. a. arnoldi*). Interestingly, all core ASVs unique to the wild *C. a. laietanus* population (i.e., ASVs 6, 29, 33, 46, 50, and 57) were also significantly more abundant in the wild population compared to the captive-bred groups of the same subspecies (Fig. 5). This pattern was not observed in *C. a. arnoldi*, where no such difference was found between the wild and captive-bred groups. Nonetheless, the same *C. a. laietanus* core ASVs listed above were also significantly more abundant in the wild *C. a. arnoldi* population when compared to its corresponding captive-bred groups.

**Fig. 5:**
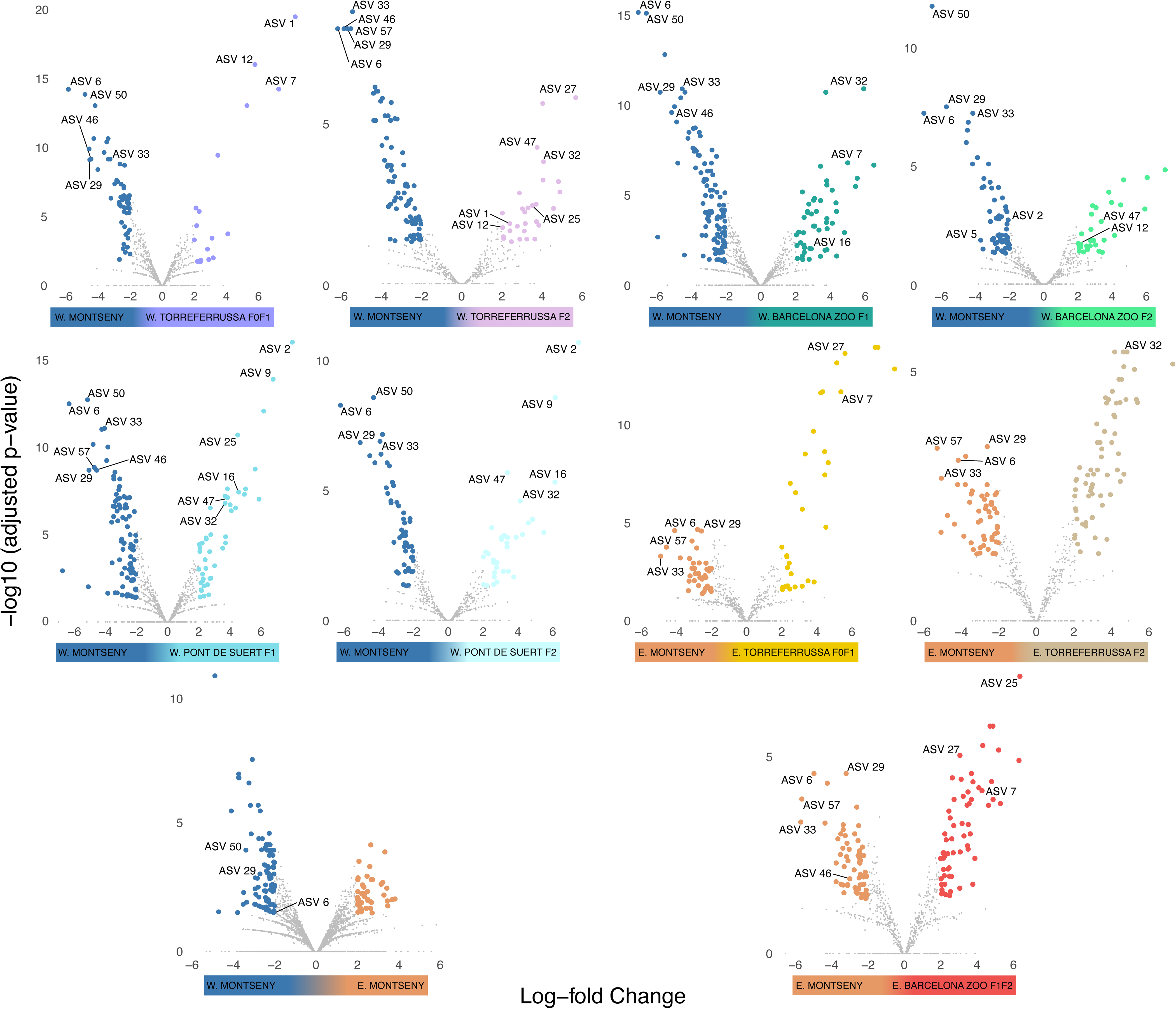
ANCOMBC analyses to test for significant differences in relative abundance between wild populations and captive-bred groups of *C. arnoldi*. The X-axis indicates the log-fold change in abundance of each ASV on a logarithmic scale, and the Y-axis indicates the statistical significance of the differences detected by doing -log_10_ (p-value). Colored dots indicate ASVs with (|Log_2_ (FC)|) > 2 and *p*-value < 0.05. W: Western Montseny brook newt (*C. a. laietanus*); E: Eastern Montseny brook newt (*C. a. arnoldi*).

Apart from the W. Torreferrussa F0F1 and F2 groups, few significant differences were detected between captive-bred groups originating from the same facility or location (<u>Fig. S2</u>).

To continue assessing the natural composition of *C. arnoldi*’s skin microbiome, we investigated putative shifts of microbiome composition at each breeding center by comparing the relative abundance of ASVs from wild populations to the captive founders (F0) and first-generation (F1) groups and then to the second generation groups (F2) (Fig. 6). From this it was revealed that the wild populations of *C. arnoldi* shared over 90% of their microbiome (<u>Fig. S3</u>), a higher proportion than any of the captive-breeding groups with their respective wild group. In both the Barcelona Zoo and Pont de Suert breeding centers, the microbiome tended to remain stable across generations (Fig. 6). However, they still lacked a major percentage of bacteria present in wild newts (such as ASV 5 and 6) and had notable differences in their relative bacteria abundances. As for the Torreferrussa breeding center, the founders and first-generation groups seem to have gone through a microbiome bottleneck, leaving few ASVs to dominate their microbiome. On the contrary, F2 generations from this center seem to be improving their microbiome’s repertoire (Fig. 6). Moreover, F2 groups from both subspecies seem to diverge in their microbiome compared to the wild populations (<u>Fig. S3</u>). Notably, the proliferation of Alphaproteobacteria in the *C. a. arnoldi* F2 groups was remarkable (Fig. 6D & E), as they do not have much presence in the wild group nor in E. Torreferrussa F0F1 group. Yet, they considerably expand in the E. Torreferrussa F2 group and in the Barcelona Zoo F1F2 group.

**Fig. 6:**
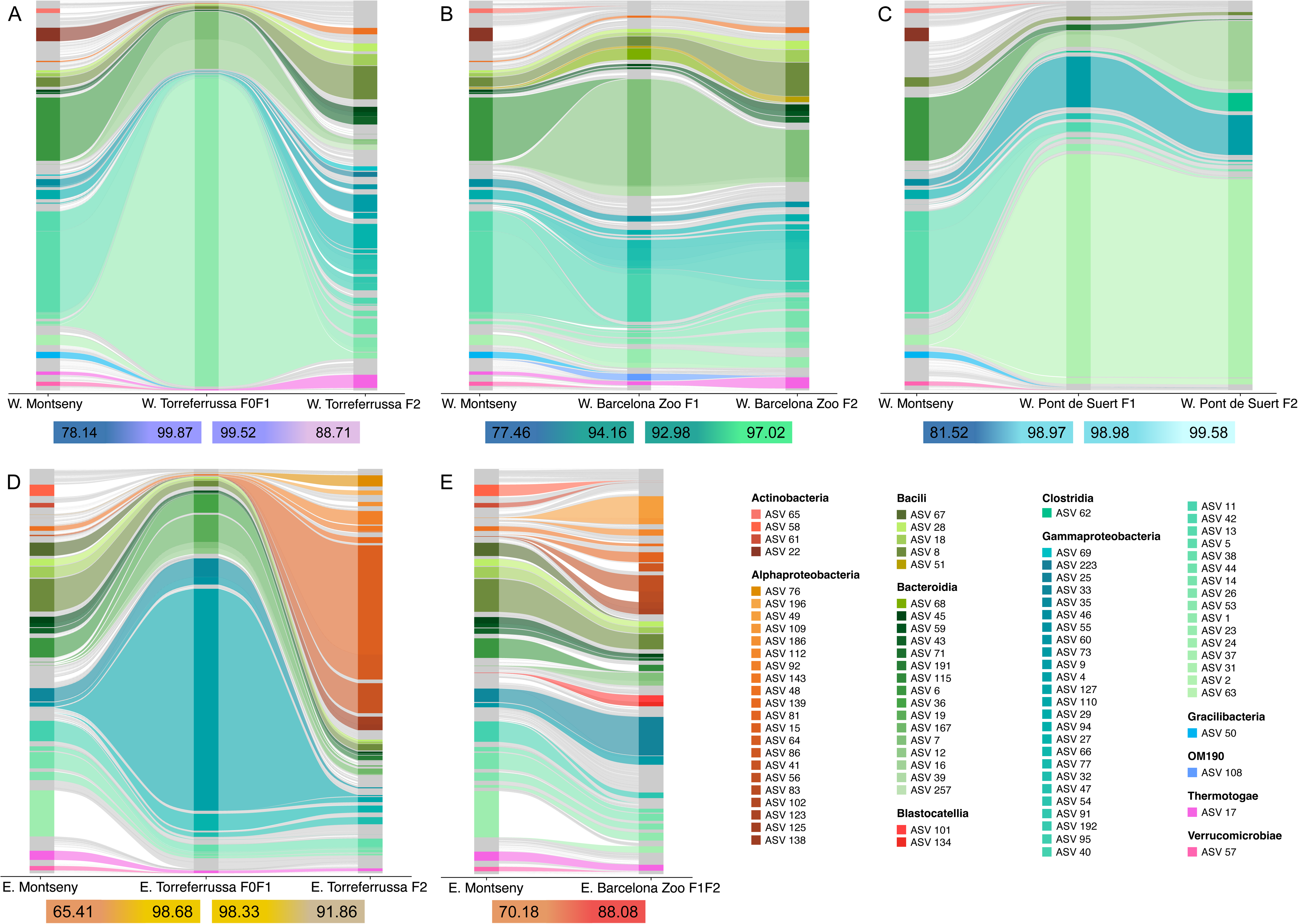
Microbiome transferability and relative abundance of shared ASVs across breeding centers of both subspecies of *C. arnoldi*. ASVs are colored according to taxonomic Class when their relative abundance equals or exceeds 1%. The percentage of total shared ASVs between groups are shown below each graph. W: Western Montseny brook newt (*C. a. laietanus*); E: Eastern Montseny brook newt (*C. a. arnoldi*).

## DISCUSSION

This study successfully described the microbiome of *C. arnoldi*, a Critically Endangered newt with a breeding program that aims to reintroduce captive-bred specimens into their natural habitat.

Results show that both *C. arnoldi* subspecies had significantly different microbiomes but had similar variance within each one. These differences could be due to their habitat, as *C. a. laietanus* tends to live in warmer streams covered by holm oaks, whereas *C. a. arnoldi* does so in streams covered by beech and other deciduous trees [19]. This could mean that each subspecies has acclimatized their skin microbiome according to or to suit their environment better, as seen in previous studies [40]. There are also core bacteria species for both subspecies, such as *Streptococcus varani*, *Oceanotoga* sp., or *Vibrio* sp., suggesting that they may be a part of the natural state of this species and that they might have an impact on the survivability of the Montseny Brook Newt.

Captive-bred groups showed lower alpha diversity values than their wild counterparts and differed in microbiome composition. This is not unexpected, as the amphibian’s microbiome cahnges are described according to sex [41], diet [42], habitat [43, 44], seasonally [45, 46] or under captivity [47], and these are clear changes between captive and wild conditions. However, since captive specimens are under much more stable and controlled conditions, it was striking that the Torreferrussa F0F1 and F2 groups of both subspecies were so different from one another (Fig. 3). It is especially surprising bearing in mind that they are under very similar environmental conditions, with low availability for a varied microbiome, and more so when considering amphibians have been associated with vertical transmission of bacteria [9]. Vertical transmission may also explain the presence of the three most abundant ASVs in the W. Pont de Suert F2 group, as they were not detected through our sampling method in the water but were very abundant in their skin microbiome. The marked difference in bacterial abundance between wild and captive-bred samples is a striking finding. Such discrepancies may have important implications for the survival of reintroduced *ex-situ* individuals, as the missing bacterial taxa could play a key role in conferring resistance to natural pathogens, including the chytrid fungi responsible for chytridiomycosis. Further research is needed to assess the functional role of *C. arnoldi*’s microbiome, particularly to understand the potential impact of pathogen exposure and the contribution of absent ASVs to the survival of captive-bred individuals following reintroduction.

An intriguing example that stands out for having a significantly lower abundance in captive samples is ASV 6, identified as *Arcicella* sp; commonly found in freshwater surface-dwelling microorganisms. Other examples of this disparity are ASV 29 and ASV 33, both *Rhodoferax* sp, a genus usually isolated from freshwater environments with putative detoxifying capacities [48]. On the other hand, certain bacteria are more abundant in captive samples than wild ones, like ASV 7, a *Flavobacterium*. This genus is widespread in water-related places and can resist water-cleansing methods like chlorine [49] and antimicrobial products [50, 51]. Some *Flavobacterium* species possess antifungal properties [52], while others have been associated with increased abundance in physiologically stressed hosts [53], in some cases reaching levels that can be lethal for certain species [54]. Similarly, ASV 1 stands out as differentially abundant in W. Torreferrussa F0F1 (Fig. 5), totally dominating this group’s microbiome. This ASV is an *Acinetobacter*, a functionally diverse genus that has been proven to have both *Bd-*inhibiting species [52] and pathogenic ones [55]. In the same way as *Flavobacterium*, *Acinetobacter* seems to thrive on amphibian skin when the host is under physiological stress [53], likely highlighting a health issue in the W. Torreferrussa F0F1 population.

A concerning issue for future reintroduction efforts is the markedly lower microbiome diversity observed in captive-bred groups compared to their wild counterparts in both subspecies. This could be, amongst many reasons, due to a lack of bacteria diversity in their environment, resulting from the change from a natural environment to an artificial one, or the water treatment as part of each center’s policy. Although the extent of microbiome plasticity in *C. arnoldi* remains unknown, exposure to their natural habitat may be sufficient to restore a microbiome similar to that of wild individuals -or at least a different but functionally effective one. In any case, providing the most realistic and ecologically relevant environment possible for captive-bred newts is essential to facilitate successful reintroduction. This should include consideration of the natural composition and diversity of the *C. arnoldi* microbiome. The importance of maintaining a diverse microbiome lies in its role in reducing the risk of potential *Bd* infections, as microbiome homogeneity has been associated with decreased survival when facing this disease [9], or other natural pathogens. Therefore, preserving or restoring microbiome diversity is a key factor in improving the reintroduction success of captive-bred *C. arnoldi* individuals into their natural environment.

One potential strategy to enhance microbiome diversity is the use of probiotic treatments, which have shown promising results in previous studies [56]. In the long term, microbiome diversity could be enhanced within captive-breeding centers by gradually introducing natural substrates and water into the tanks -provided they are first screened for common amphibian pathogens. In the short term, and to avoid disrupting the current captive conditions, a soft-release strategy may be more effective. This approach involves temporarily confining individuals at the release site to allow acclimatization before full release [57], and has been shown to yield positive outcomes in some amphibian species [58].

This is particularly important when considering that amphibians can horizontally transmit skin bacteria [40]. In such cases, all individuals undergoing the soft-release procedure would eventually share the acquired bacteria upon reintroduction to the same location. Given the species’ microbiome plasticity, there is promise that reintroduced specimens can adapt their microbiome to withstand potential challenges better. Furthermore, the captive-bred *C. arnoldi* groups appeared to acquire different environmental bacteria compared to their wild counterparts, yet lacked several core ASVs that may play critical roles in their natural habitat. These missing ASVs should be prioritized in the development of probiotic treatments. In their absence, the microbiomes of captive individuals were often dominated by bacterial taxa not found in wild specimens. For instance, *Flectobacillus fontis* (ASV 18) was prevalent in captive-bred *C. a. arnoldi* groups, while *Flavobacterium* sp. (ASV 7) dominated the microbiome of captive-bred *C. a. laietanus* groups. Both taxa have also been detected in other captive amphibian species [40, 59], suggesting they may represent a microbial signature of captivity.

Moreover, some groups showed noticeable differences between the relative abundance of certain ASVs in their skin microbiome and in the surrounding water. For instance, individuals from the W. Pont de Suert F1 group exhibited high relative abundances of *Caenimonas* sp. on their skin, despite this ASV ranking 102^nd^ out of 144 detected ASVs in their aquatic environment (<u>Table S3</u>). Specimens from the E. Barcelona Zoo F1F2 group also illustrated this mismatch, with ASV 25 being relatively abundant in their skin microbiomes despite ranking as the 145^th^ most abundant ASV in their surrounding water (<u>Table S3</u>). Although the idea that amphibians may selectively recruit rare environmental bacteria is well established [60], it remains intriguing that individuals preferentially acquired these low-abundance ASVs over others such as *Erwinia* sp. or *Arcinella* sp., which were more abundant in the water and also commonly found in the wild populations’ skin microbiomes. The criteria for determining which bacteria become part of an individual’s microbiome remain poorly understood, but are likely influenced by a combination of host-specific traits, bacterial characteristics, host-microbe interactions, and environmental conditions. Given that amphibians may harbor bacterial communities particularly suited to their environmental and physiological needs and that these bacteria can be transmitted between individuals, further exploring phylosymbiosis in *C. arnoldi* would be of great interest. This concept describes the coevolution of the species and their microbiomes in addition to environmental effects, tracing parallelisms between the microbial community and the host species. Therefore, it would mean that an amphibian microbial community can influence host evolution through composition and functional effects [61], which can in turn affect the species’ ecology, physiology and behavior.

This study builds on previous work examining the microbiomes of wild and captive amphibians by providing the detailed characterization of the microbiome composition of *Calotriton arnoldi*, one of Europe’s most threatened amphibians. Our findings highlight the importance of maintaining a healthy and diverse microbiome when this species is kept outside its natural habitat. Notably, the results also reveal *C. arnoldi*’s microbiome’s plasticity, as each captive-bred group developed significantly different microbiomes. While much remains to be understood, our findings raise important questions regarding reintroduction success. For instance, it is critical to assess whether microbiome composition is directly linked to survival after reintroduction, particularly in the face of pathogenic threats, or how the microbiome of released individuals will evolve. Will it reflect the community acquired in captivity, shift toward that of wild populations of the same subspecies, or develop into a distinct new assemblage? Moreover, establishing a baseline for the presence or absence of key bacterial taxa will be essential for designing future probiotic treatments, especially given the microbiome’s immunological role and the presence of *Bd* and *Bsal* in the region. In conclusion, a more comprehensive understanding of *C. arnoldi*’s microbiome and its ecological significance is vital for informing effective conservation strategies and ensuring the success of reintroduction programs.

## ACKNOWLEDGMENTS

This work was supported by the Planetary Wellbeing Initiative research actions (2021) PLAWB00621 project “Identification and isolation of probiotic bacteria to protect the Critically Endangered Montseny Brook Newt from chytridiomycosis” funded by Universitat Pompeu Fabra and awarded to Elena Bosch, Javier del Campo, and Salvador Carranza. Sergi A. Tulloch Jiménez was funded by the JAE Intro fellowship JAEINT_22_02190, awarded by the Spanish Foundation for Science and Technology (FECYT) under the research programs of the Junta de Ampliación de Estudios. Maria Estarellas was funded by an FPI grant from the Ministerio de Ciencia, Innovación y Universidades, Spain (PRE2022-101473). The LIFE project (LIFE15 NAT/SE/000757) financially supported some authors of this work.

## DATA AVAILABILITY

The raw reads for the project have been deposited on NCBI SRA (BioProject: PRJNA1265703). Code used for analysis is available upon request from the corresponding author. Taxonomy table, statystical results, relative abundances, core ASVs and one-to-one ANCOM results are available with the Supplementary Material.

## SUPPLEMENTARY MATERIAL

**Fig. S1: Microbiome composition and variance differences between *C. arnoldi* and their environmental water. A** Aitchison Distance PCA with 95% confidence interval ellipse for *C. arnoldi* and water samples. **B** Bubble plot representing Genera with >1% relative abundance in at least one of the groups for all groups of *C. arnoldi* and their respective water sample, with Genera color-coded according to taxonomic Class. **C** Heatmap representing the most abundant ASVs across groups and water samples, with ASVs color-coded according to taxonomic Class. W: Western Montseny brook newt (*C. a. laietanus*); E: Eastern Montseny brook newt (*C. a. arnoldi*).

**Fig. S2: ANCOMBC analyses to test for significant differences in relative abundance between captive-bred groups of the same location for both subspecies of *C. arnoldi*.** The X-axis indicates each ASV’s log-fold change in abundance, and the Y-axis indicates the statistical significance of the differences detected by doing -log10 (p-value). W: Western Montseny brook newt (*C. a*. *laietanus*); E: Eastern Montseny brook newt (*C. a*. *arnoldi*).

**Fig. S3: Microbiome transferability and relative abundance of shared ASVs between wild populations and F2 captive-bred groups for both subspecies of *C. arnoldi.*** ASVs are coloured according to taxonomic Class when their relative abundance equals or exceeds 1%. The percentage of total shared ASVs between groups are shown under each graph. W: Western Montseny brook newt (*C. a. laietanus*); E: Eastern Montseny brook newt (*C. a. arnoldi*).

**Table S1: Taxonomic classification of ASVs.**

**Table S2: Linear model results and subsequent pairwise analyses for composition and variance of each subspecies of *C. arnoldi*.** Sheet 1 has all analyses related to *C. a. laietanus*. Sheet 2 has all analyses related to *C. a. arnoldi*. W: Western Montseny brook newt (*C. a. laietanus*); E: Eastern Montseny brook newt (*C. a. arnoldi*).

**Table S3: Relative abundances of each ASV for different categorizations of the *C. arnoldi* and water samples.** Sheet 1 has ASV relative abundances only for wild samples, including *C. a. laietanus* samples and *C. a. arnoldi* samples and both wild groups merged together. Sheet 2 has ASV relative abundances for only captive *C. a. laietanus* samples, including all the respective groups and all merged captive *C. a. laietanus* samples. Sheet 3 has ASV relative abundances for only captive *C. a. arnoldi* samples, including all the respective groups and all captive *C. a. arnoldi* samples merged. Sheet 4 includes ASV relative abundances for all water samples. W: Western Montseny brook newt (*C. a. laietanus*); E: Eastern Montseny brook newt (*C. a. arnoldi*).

**Table S4: Core ASVs for each wild population and captive-bred group of *C. arnoldi*.** Core ASVs were defined as those shared by at least 75% of the samples with a relative abundance equal or greater than 0.1%. W: Western Montseny brook newt (*C. a. laietanus*); E: Eastern Montseny brook newt (*C. a. arnoldi*).

**Table S5: ANCOMBC analyses for different pairs of *C. arnoldi* groups.** The full output of the results of each ANCOMBC performed and illustrated in Fig. 5 and Fig. S2. Each Sheet corresponds to a compared pair of groups in the following order: Sheet 1: W. Montseny vs W. Torreferrussa F0F1; Sheet 2: W. Montseny vs W. Torreferrussa F2; Sheet 3: W. Torreferrussa F0F1 vs W. Torreferussa F2; Sheet 4: W. Montseny vs W. Pont de Suert F1; Sheet 5: W. Montseny vs W. Pont de Suert F2; Sheet 6: W. Pont de Suert F1 vs W. Pont de Suert F2; Sheet 7: W. Montseny vs W. Barcelona Zoo F1; Sheet 8: W. Montseny vs W. Barcelona Zoo F2; Sheet 9: W. Barcelona Zoo F1 vs W. Barcelona Zoo F2; Sheet 10: W. Montseny vs E. Montseny; Sheet 11: E. Montseny vs E. Torreferrussa F0F1; Sheet 12: E. Torreferrussa F2; Sheet 13: E. Torreferrussa F0F1 vs E. Torreferrussa F2; Sheet 14: E. Montseny vs E. Barcelona Zoo F1F2. W: Western Montseny brook newt (*C. a. laietanus*); E: Eastern Montseny brook newt (*C. a. arnoldi*).

