## Supplementary figures and images for "Comparative microbiome analyses reveal differences between wild populations and captive groups of the Montseny Brook Newt (*Calotriton arnoldi*)"

### Fig_S1

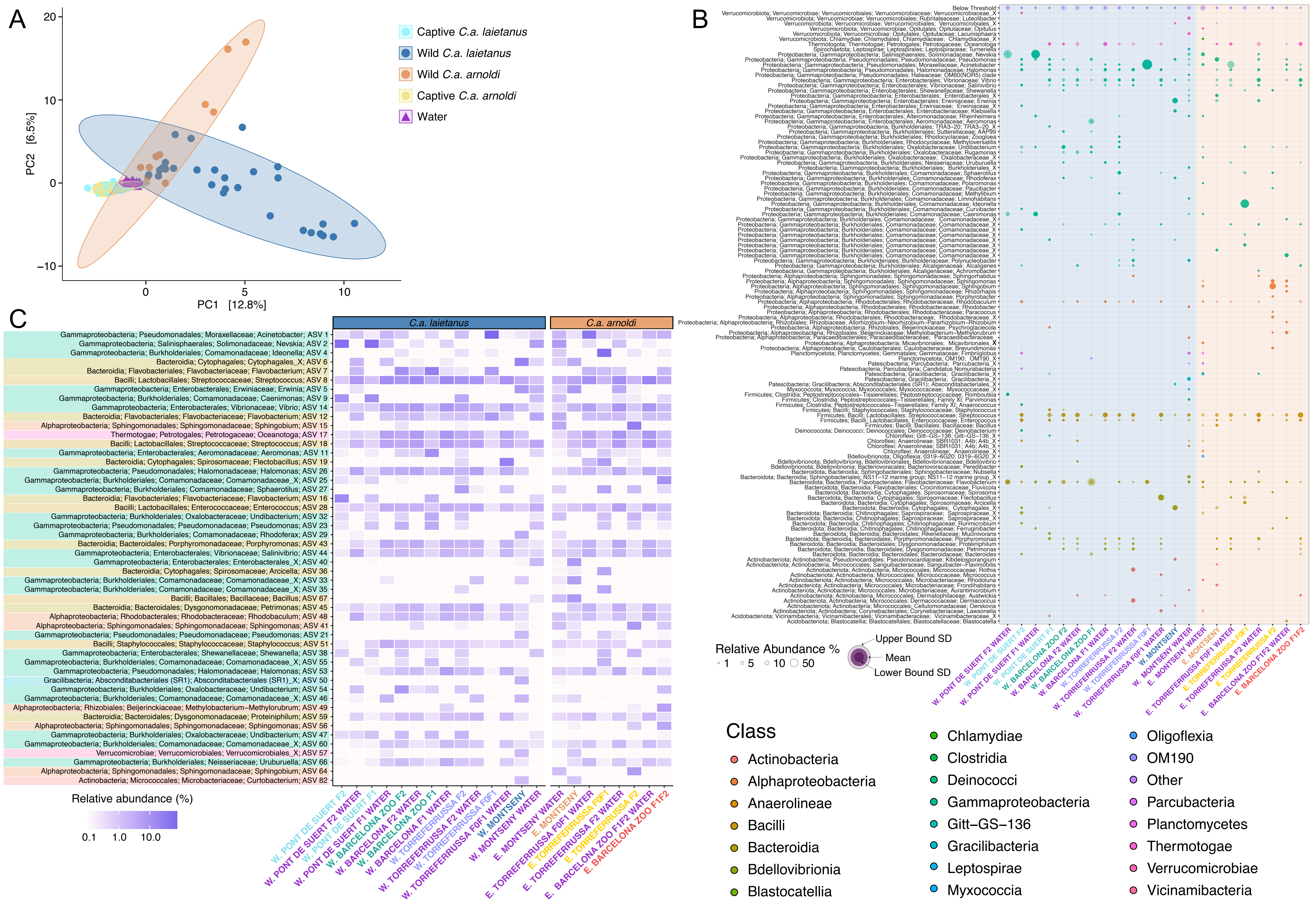

### Supplementary Figure 2

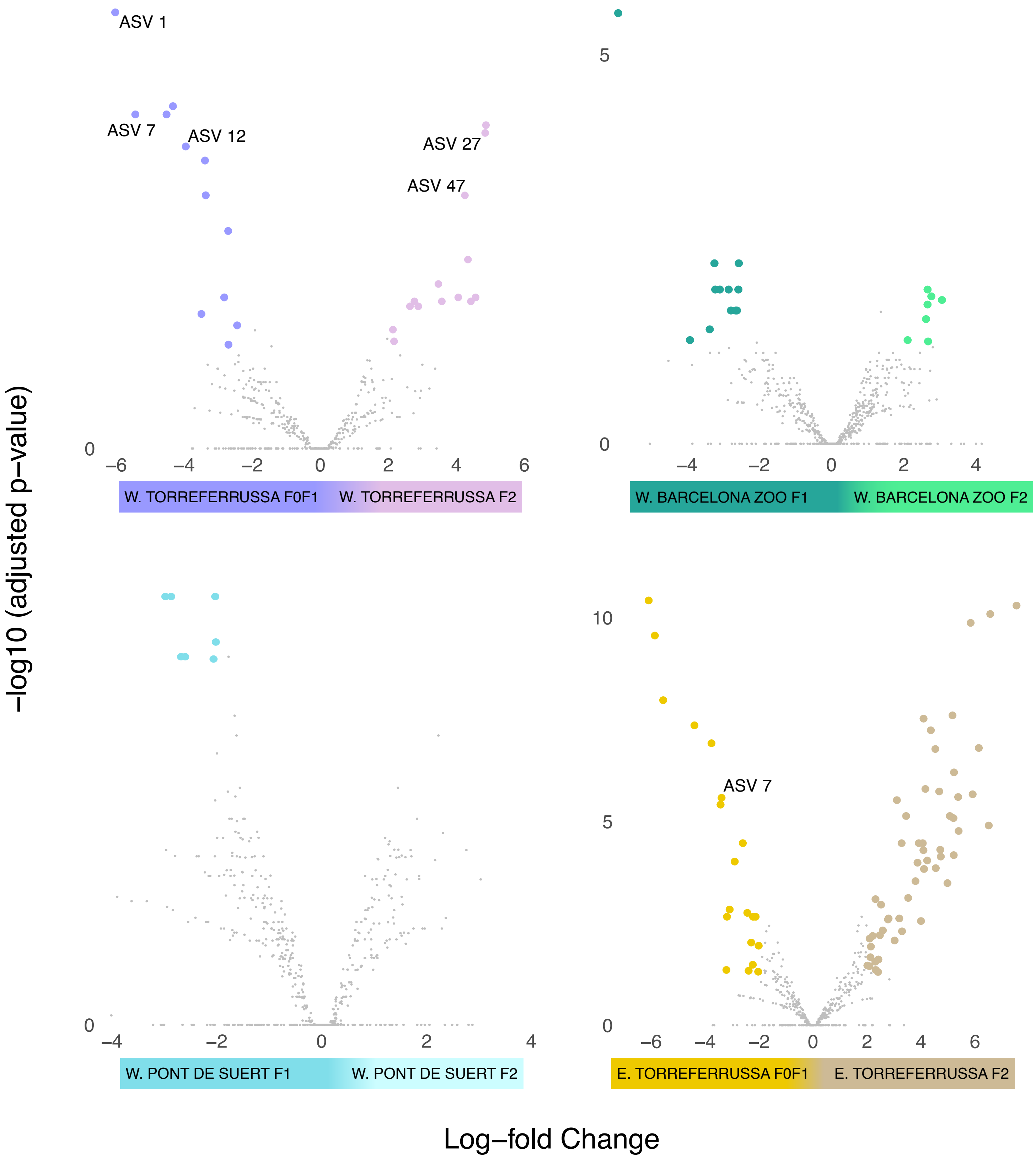

### Supplementary Figure 3

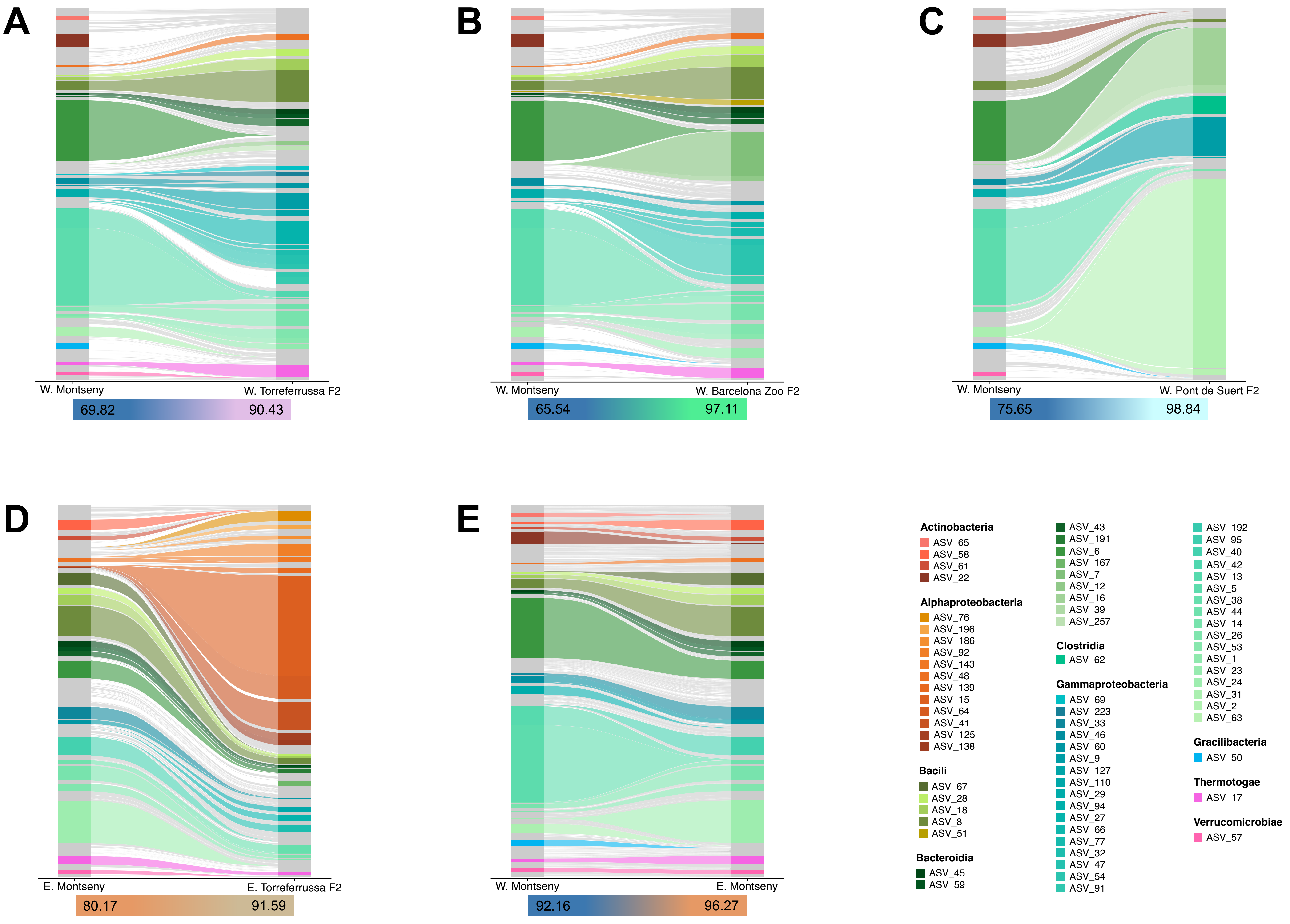
